# Open access in silico tools to predict the ADMET profiling and PASS (Prediction of Activity Spectra for Substances of Bioactive compounds of Garlic (Allium sativum L.)

**DOI:** 10.1101/2021.07.18.452815

**Authors:** Ivan Vito Ferrari

**Affiliations:** Department of Industrial Engineering and LIME Laboratory, University of Rome Tor Vergata, Via del Politecnico 1, 00133 Rome, Italy

**Keywords:** in silico, ADMET, drug, open access, prediction, PASS Online ((Prediction of Activity Spectra for Substances), Garlic Compounds, S-allylcysteine ( SAC) and S-allylmercaptocysteine (SAMC)

## Abstract

**Background:** Garlic (Allium sativum L.) is a common spice with many health benefits, mainly due to its diverse bioactive compounds, (see below) such as organic sulphides, saponins, phenolic compounds, and polysaccharides. Several studies have demonstrated its functions such as anti-inflammatory, antibacterial, and antiviral, antioxidant, cardiovascular protective and anticancer property. In this work we have investigated the main bioactive components of garlic through a bioinformatics approach. Indeed, we are in an era of bioinformatics where we can predict data in the fields of medicine. Approaches with open access in silico tools have revolutionized disease management due to early prediction of the absorption, distribution, metabolism, excretion, and toxicity (ADMET) profiles of the chemically designed and eco-friendly next-generation drugs.

**Methods:** This paper encompasses the fundamental functions of open access in silico prediction tools, as PASS database (Prediction of Activity Spectra for Substances) that it estimates the probable biological activity profiles for compounds. This paper also aims to help support new researchers in the field of drug design and to investigate best bioactive compounds in garlic.

**Results:** screening through each of pharmacokinetic criteria resulted in identification of Garlic compounds that adhere to all the ADMET properties.

**Conclusions:** It was established an open-access database (PASS database, available bioinformatics tool SwissADME, PreADMET pkCSM database) servers were employed to determine the ADMET (metabolism, distribution, excretion, absorption, and toxicity) attributes of garlic molecules and to enable identification of promising molecules that follow ADMET properties.

**Graphical abstract:** 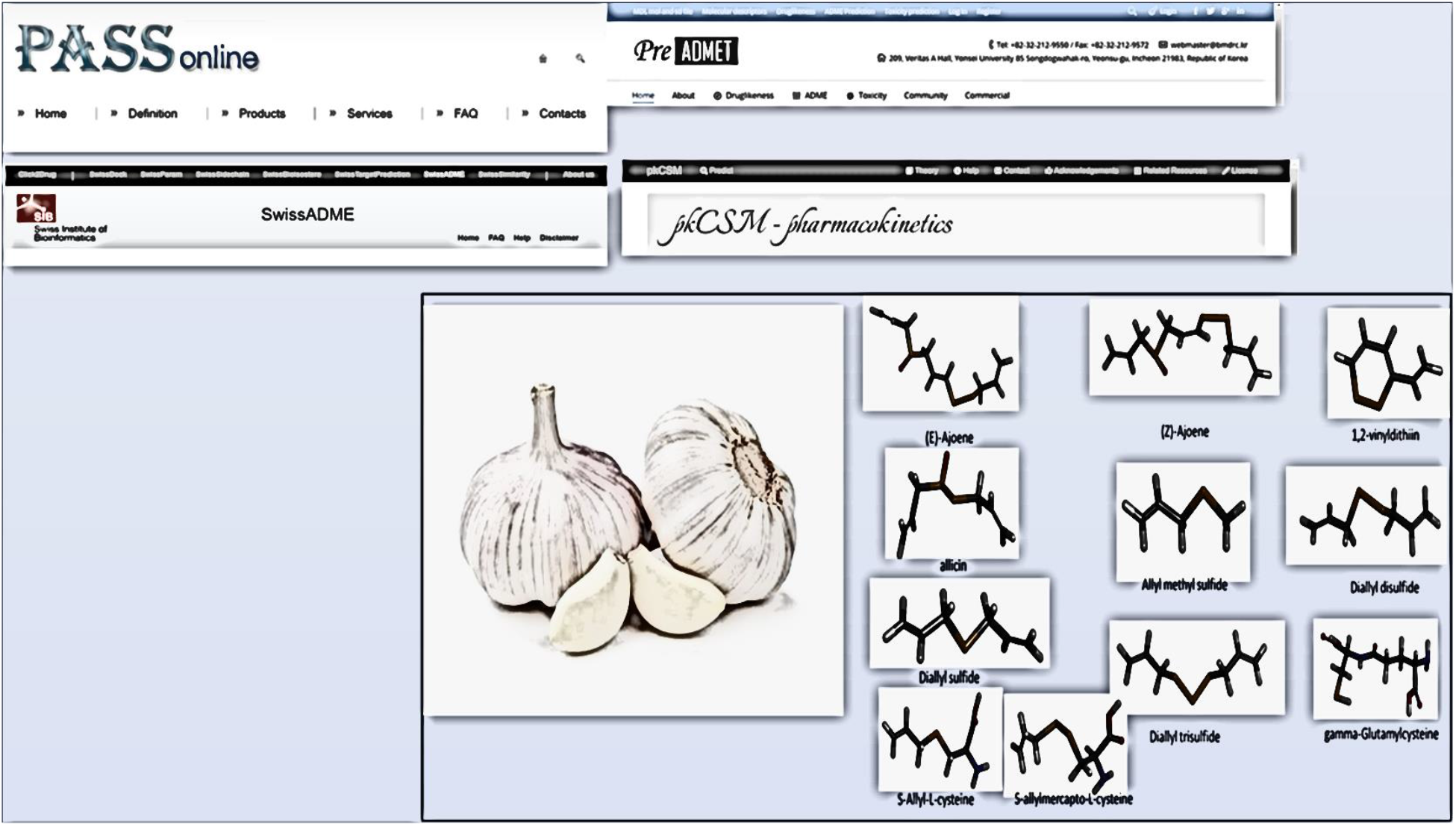

## 1. Introduction

Garlic (Allium sativum L.) is a species in the onion genus Allium. This is a common spice with many health benefits, mainly due to its diverse bioactive compounds, (see below fig.1) such as organic sulphides, saponins, phenolic compounds, and polysaccharides. Garlic has been demonstrated to exhibit potentially beneficial for cancer prevention. Several studies have demonstrated its functions such as anti-inflammatory, antibacterial, and antiviral, antioxidant, cardiovascular protective. anticancer property. [1–2] Observations over the past years have shown that the consumption of garlic in the diet provides strong protection against cancer risk. [2]. In literature we can find some papers, where it was demonstrated decreased rates stomach cancer associated with garlic intake. [3–5]

**Fig. 1.**
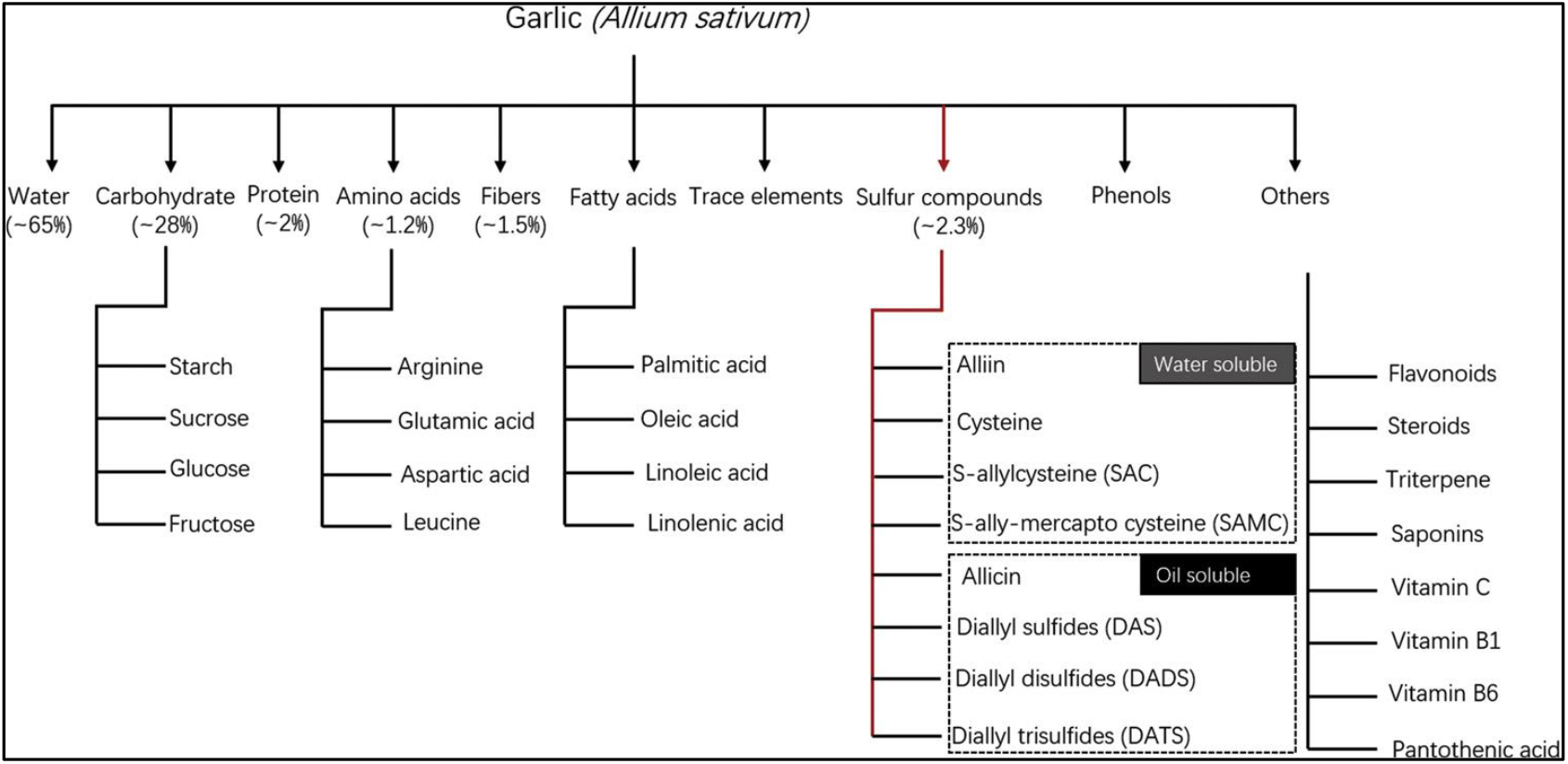
Major classification of the bioactive constituents in garlic. Generally, garlic bulb contains approximately 65 % water, 28 % carbohydrates (mainly fructans), 2 % protein (mainly alliin), 1.2 % free amino acids (mainly arginine), 1.5 % fiber, and 2.3 %organosulfur compounds [2]

In this work we have investigated the main bioactive components of garlic through a bioinformatics approach. Indeed, we are in an era of bioinformatics where we can predict data in the fields of medicine. Approaches with open access in silico tools have revolutionized disease management due to early prediction of the absorption, distribution, metabolism, excretion, and toxicity (ADMET) profiles of the chemically designed and eco-friendly next-generation drugs. [6–7] This paper encompasses the fundamental functions of open access in silico prediction tools, as PASS database (Prediction of Activity Spectra for Substances) that it estimates the probable biological activity profiles for compounds. [8–9] This paper also aims to help support the researchers in the field of drug design and to investigate best bioactive compounds in garlic. As it has been before, Garlic contains 0.1-0.36% of a volatile oil these volatile compounds are generally considered to be responsible for most of the pharmacological properties of garlic. Garlic contains at least 33 sulfur compounds like aliin, allicin, ajoene, allylpropl, diallyl, trisulfide, s-allylcysteine, vinyldithiines, S-allylmercaptocystein, and others. Particular attention has been given to sulphide compounds of garlic for their anti-tumour properties, S-allylcysteine ( SAC) and S-allylmercaptocysteine (SAMC) ( See below fig.1–2) [10–20] It was established an open-access database (PASS database, available bioinformatics tool SwissADME, PreADMET pkCSM database) servers were employed to determine the ADMET (metabolism, distribution, excretion, absorption, and toxicity) attributes of garlic molecules and to enable identification of promising molecules that follow ADMET properties. [6–9]

**Fig 2.**
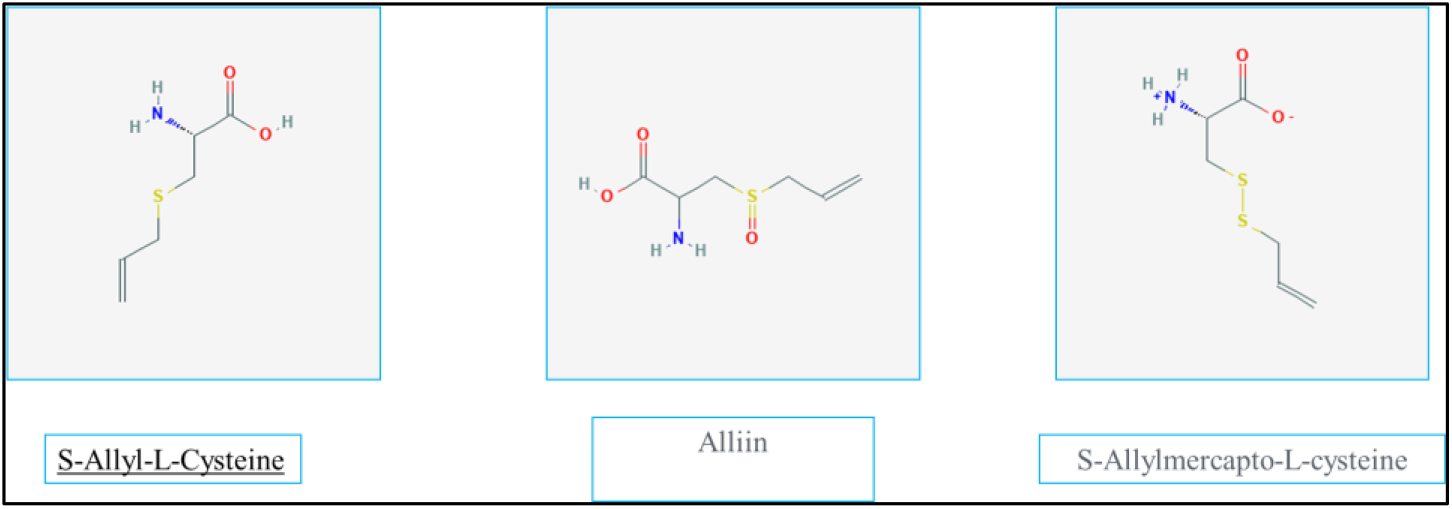
bioactive constituents in garlic sulphur compounds

## 2. Materials and methods

This paper encompasses the fundamental functions of open access in silico prediction tools, as PASS database, PreADMET and pkCSM (http://biosig.unimelb.edu.au/pkcsm/) servers were employed to determine the ADMET (metabolism, distribution, excretion, absorption, and toxicity) attributes of target molecules. PASS (Prediction of Activity Spectra for Substances) Online predicts over 4000 kinds of biological activity, including pharmacological effects, mechanisms of action, toxic and adverse effects, interaction with metabolic enzymes and transporters, influence on gene expression, etc. The first strategy is based on the suggestion that the more kinds of activity are predicted as probable for a compound, the more probable to find any useful pharmacological action in it. Prediction is based on the analysis of structure activity-relationships for more than 250,000 biologically active substances including drugs and drug-candidates. [8] The available bioinformatics tool SwissADME (http://www.swissadme.ch/index.php) [7] was used for finding drug-likeness attributes. Lipinski’s rule of five [6] was used to analyze the properties such as; hydrogen bond donor (HBD), hydrogen bond acceptor (HBA), molecular weight (MW), and lipophilicity (log P). PreADMET (https://preadmet.bmdrc.kr/) and pkCSM (http://biosig.unimelb.edu.au/pkcsm/) servers were employed to determine the ADMET (metabolism, distribution, excretion, absorption, and toxicity) attributes of target molecules.

### 2.1 Classification of the bioactive constituents in garlic

3D Conformer of several bioactive constituents in garlic were download from PubChem (https://pubchem.ncbi.nlm.nih.gov/) and create by Discovery Studio Biovia Visualizer Software [21] ( see fig.3)

**Fig 3.**
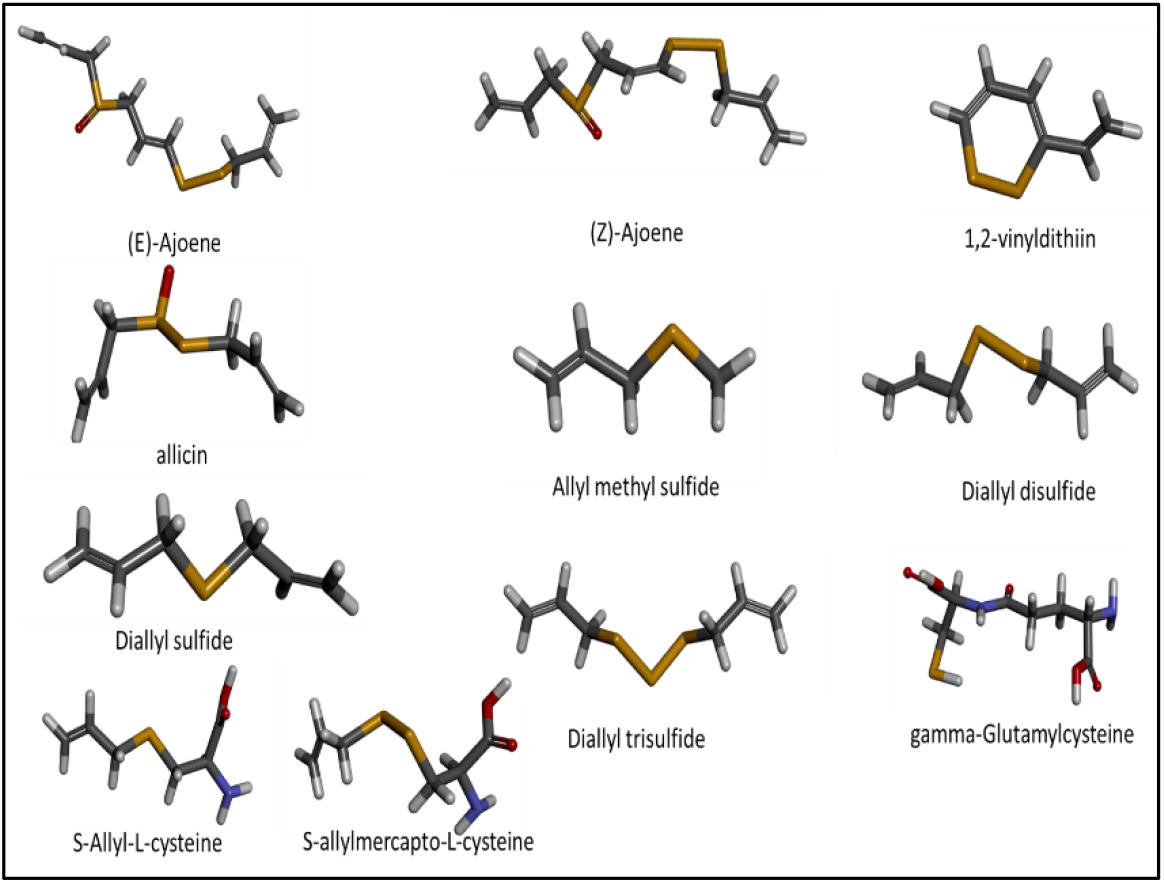
Principal Organosulfur Compounds From Garlic. Reproduced by Discovery Studio Biovia Visualizer Software [21]

## 3. Results and discussion

### 3.1 PASS database (Prediction of Activity Spectra for Substances)

PASS Online predicts over 3500 kinds of biological activity, including pharmacological effects, mechanisms of action, toxic and adverse effects, interaction with metabolic enzymes and transporters, influence on gene expression, etc. Prediction is based on the analysis of structure activity-relationships for more than 250,000 biologically active substances including drugs, drug-candidates, leads and toxic compounds. The concept of the biological activity spectrum was introduced to describe the properties of biologically active substances. The PASS (prediction of activity spectra for substances) software product, which predicts more than 300 pharmacological effects and biochemical mechanisms on the basis of the structural formula of a substance, may be efficiently used to find new targets (mechanisms) for some ligands and, conversely, to reveal new ligands for some biological targets. Average accuracy of prediction estimated in leave-one-out cross-validation procedure (each compound is excluded from the training set and its activity predicted based on SAR model obtained on the rest part of the training set) for the whole PASS training set is about 95% (Filimonov and Poroikov, 2008) [22] . Since PASS service is used by medicinal chemists, pharmacologists and toxicologists for several years (Lagunin et al., 2000) [23], there are many publications where PASS predictions were confirmed by subsequent synthesis and biological testing. To provide more accurate predictions for compounds belonging to new chemical classes and to extend the predictable area onto new biological activities, we are permanently working on enlargement of PASS training set. Input data represents a structural formula of a compound in MOL file format. The output file represents a list of activities with two probabilities Pa (probability to be active) and Pi (probability to be inactive). Pa (probability “to be active”) estimates the chance that the studied compound is belonging to the sub-class of active compounds (resembles the structures of molecules, which are the most typical in a sub-set of “actives” in PASS training set). Pi (probability “to be inactive”) estimates the chance that the studied compound is belonging to the sub-class of inactive compounds (resembles the structures of molecules, which are the most typical in a sub-set of “inactives” in PASS training set). The first strategy is based on the suggestion that the more kinds of activity are predicted as probable for a compound, the more probable to find any useful pharmacological action in it. For each compound from available set of samples the following value can be calculated: [8–9] ; [22–23]

**Fig 4.**
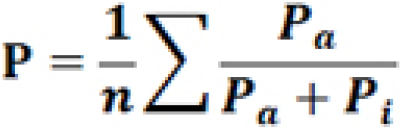
Prediction of Activity Spectra for Substances calculated : the output file represents a list of activities with two probabilities Pa (probability to be active) and Pi (probability to be inactive [8–9] ; [22–23]

In Tab 1 we report chemical-physical properties of Principal Organosulfur Compounds , S-allylcysteine ( SAC) and S-allylmercaptocysteine (SAMC) in Garlic investigated by Pass Online Server ( Prediction of Activity Spectra for Substances ) that it estimates the probable biological activity profiles for compounds. As we can see from the table 1, SAMC and SAC demonstrated to a suppressive agent against several tumours and they have several functions such as anti-inflammatory, antibacterial, and antiviral, antioxidant, cardiovascular protective. [10–20] S-allylcysteine ( SAC) and S-allylmercaptocysteine (SAMC) have a high value of 0.96-0.98 Pa (probability to be active) in human flavin-containing monooxygenase 3 (FMO3) and it has impact on enzyme activity, drug metabolism and disease. [28–29] Indeed, A flavin-containing monooxygenase (FMO) produced by A. sativum (AsFMO) was previously proposed to oxidize S-allyl-L-cysteine (SAC) to alliin, an allicin precursor. 30. Ferreira F, et all., (2013) have investigated the activity of the human flavin-containing monooxygenase (FMO) has been proposed to be impact on enzyme activity, drug metabolism and disease, like Trimethylaminuria (TMAu) or “fish odor syndrome” is a metabolic disorder characterized by the inability to convert malodorous dietarily-derived trimethylamine (TMA) to odourless TMA N-oxide by the flavin-containing monooxygenase 3 (FMO3). [30]

**Tab 1.** Prediction of Activity Spectra for Substances of Principal Organosulfur Compounds, as S-allylcysteine ( SAC) and S-allylmercaptocysteine (SAMC)

| S-Allyl-L-cysteine | C<sub>6</sub>H<sub>11</sub>NO<sub>2</sub>S | 161.22 | Pa | Pi | Activity |
| --- | --- | --- | --- | --- | --- |
|  |  |  | 0,965 | 0,002 | Flavin-containing monooxygenase substrate |
|  |  |  | 0,963 | 0,003 | CYP2E1 substrate |
|  |  |  | 0,962 | 0,003 | CYP2E substrate |
|  |  |  | 0,942 | 0,001 | S-alkylcysteine lyase inhibitor |
|  |  |  | 0,939 | 0,002 | NADPH peroxidase inhibitor |
|  |  |  | 0,933 | 0,002 | FMO3 substrate |
|  |  |  | 0,928 | 0,002 | Lysine 2,3-aminomutase inhibitor |
|  |  |  | 0,924 | 0,003 | Acylcarnitine hydrolase inhibitor |
|  |  |  | 0,905 | 0,004 | Arylacetonitrilase inhibitor |
| S-allylmercapto-L-cysteine | C<sub>6</sub>H<sub>11</sub>NO<sub>2</sub>S<sub>2</sub> | 193.3 | Pa | Pi | Activity |
|  |  |  | 0,974 | 0,002 | Flavin-containing monooxygenase substrate |
|  |  |  | 0,967 | 0,001 | FMO3 substrate |
|  |  |  | 0,939 | 0,002 | NADPH peroxidase inhibitor |
|  |  |  | 0,939 | 0,002 | Protein-disulfide reductase (glutathione) inhibitor |
|  |  |  | 0,934 | 0,003 | CYP2E1 substrate |
|  |  |  | 0,933 | 0,003 | CYP2E substrate |
|  |  |  | 0,932 | 0,003 | Antiseborrheic |
|  |  |  | 0,928 | 0,003 | Lipid metabolism regulator |
|  |  |  | 0,925 | 0,004 | Apoptosis agonist |
|  |  |  | 0,924 | 0,003 | Acylcarnitine hydrolase inhibitor |
|  |  |  | 0,919 | 0,001 | Cysteamine dioxygenase inhibitor |
|  |  |  | 0,916 | 0,003 | Arylacetonitrilase inhibitor |
|  |  |  | 0,909 | 0,002 | S-alkylcysteine lyase inhibitor |
|  |  |  | 0,903 | 0,005 | TP53 expression enhancer |

### 3.2 In silico Organosulfur Compounds analysis and ADMET profiling

The available bioinformatics tool SwissADME (http://www.swissadme.ch/index.php) [7,24] was used for finding drug-likeness attributes. Lipinski’s rule of five [6] was used to analyze the properties such as; hydrogen bond donor (HBD), hydrogen bond acceptor (HBA), molecular weight (MW), and lipophilicity (log P). PreADMET (https://preadmet.bmdrc.kr/) and pkCSM (http://biosig.unimelb.edu.au/pkcsm/) servers were employed to determine the ADMET (metabolism, distribution, excretion, absorption, and toxicity) attributes of target molecules.

#### 3.2.1 SwissADME (http://www.swissadme.ch/index.php)

This website allows you to compute physicochemical descriptors as well as to predict ADME parameters, pharmacokinetic properties, druglike nature and medicinal chemistry friendliness of one or multiple small molecules to support drug discovery. This web service is a free web tool to evaluate pharmacokinetics, drug-likeness and medicinal chemistry friendliness of small molecules.[24]

**Fig 5.**
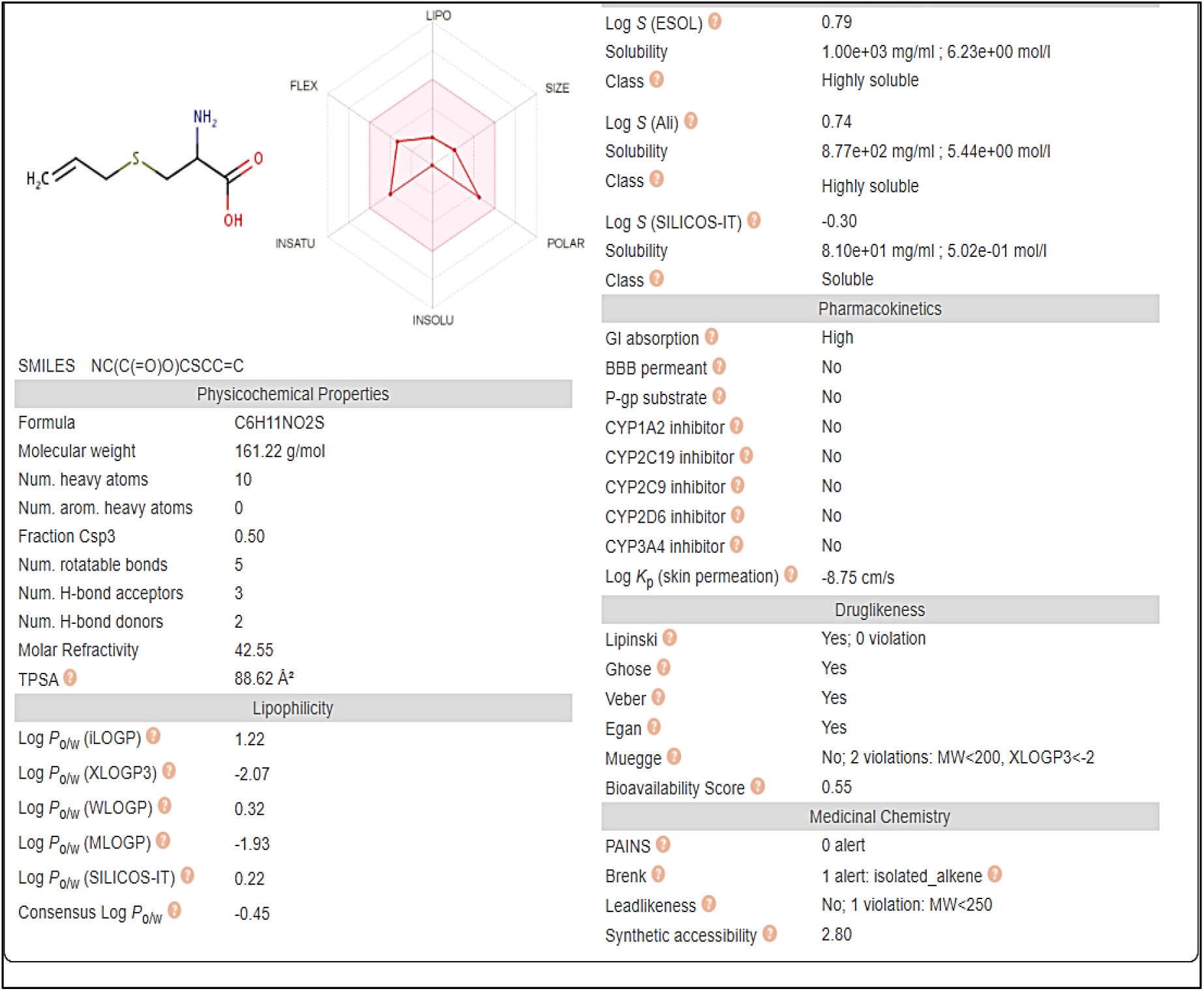
physicochemical descriptor, ADME parameters, pharmacokinetic properties, druglike nature and medicinal chemistry friendliness of S-allylcysteine ( SAC ) predicted by SwissADME Database SwissADME (http://www.swissadme.ch/index.php)

**Fig 6.**
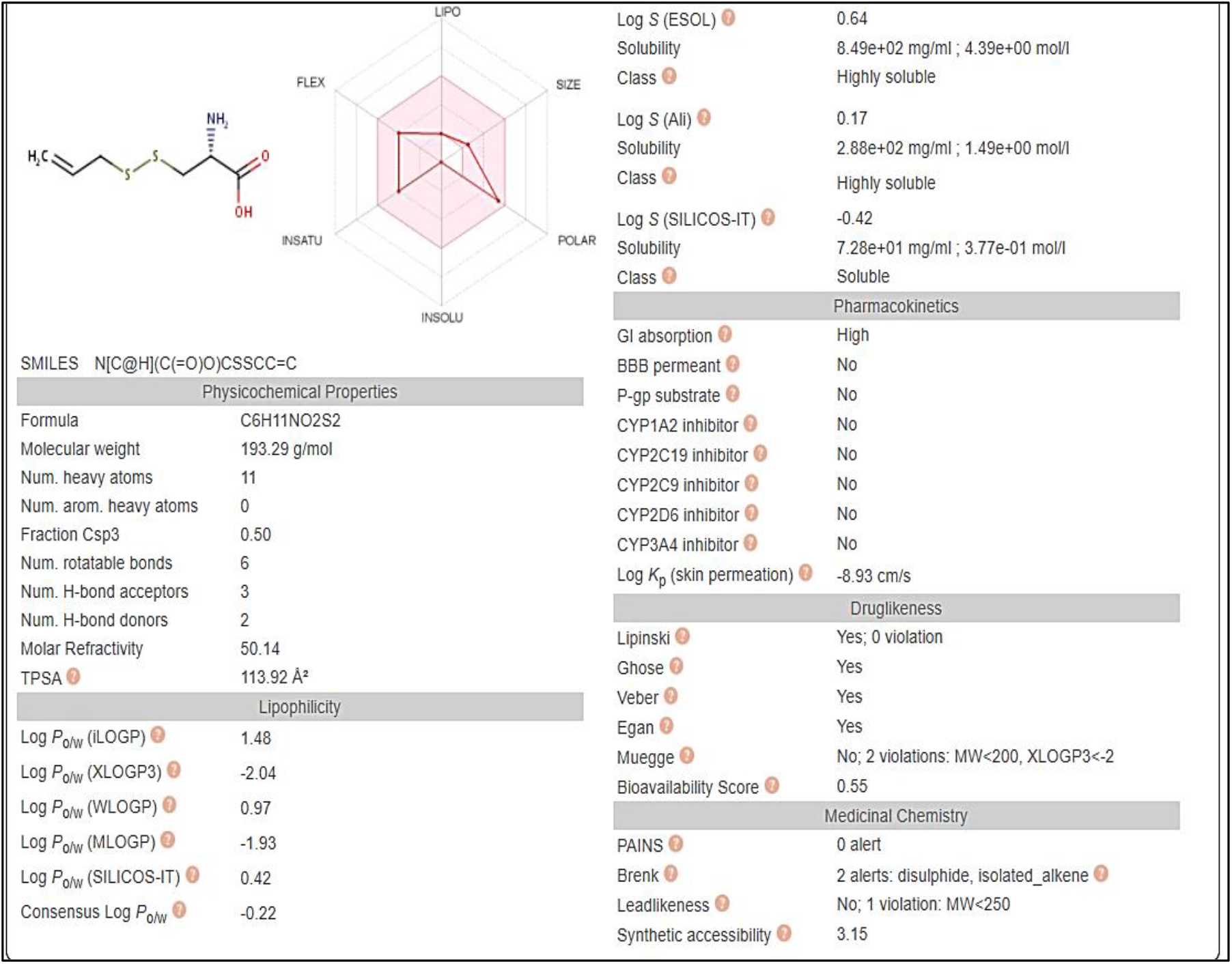
physicochemical descriptors), ADME parameters, pharmacokinetic properties, druglike nature and medicinal chemistry friendliness of S-allylmercaptocysteine (SAMC) predicted by SwissADME Database SwissADME (http://www.swissadme.ch/index.php)

#### 3.2.2 PreADMET (https://preadmet.bmdrc.kr/)

PreADMET is a web-based application for predicting ADME data and building drug-like library using in silico method. It was describe a new web-based application called PreADMET, which has been developed in response to a need for rapid prediction of drug-likeness and ADME/Tox data. [25–26] In figure 7 we report ADMET (metabolism, distribution, excretion, absorption, and toxicity) attributes of S-allylcysteine ( SAC ) ) predicted by PreADMET (https://preadmet.bmdrc.kr/ ):

- BBB (in vivo blood-brain barrier penetration (C.brain/C.blood) ≫ 0.229899 Value
- Calculated water solubility value in buffer system by SK atomic types (mg/L) ≫ 92985.4 Value
- in vitro Caco2 cell permeability (Human colorectal carcinoma; nm/sec) ≫ 6.65342 Value
- in vitro Cytochrome P450 2C19 inhibition ≫ Inhibitor
- in vitro Cytochrome P450 2C9 inhibition ≫ Inhibitor
- in vitro Cytochrome P450 2D6 inhibition ≫ Inhibitor
- in vitro Cytochrome P450 2D6 substrate ≫ No value
- in vitro Cytochrome P450 3A4 inhibition ≫ No value
- Human intestinal absorption (HIA, %) ≫ 81.972219 Value
- in vitro MDCK cell permeability (Mandin Darby Canine Kidney) ≫ 209.694
- in vitro P-glecoprotein inhibition≫ No value
- in vitro plasma protein binding (%) ≫ 11.674627
- in vitro skin permeability (transdermal delivery) ≫ −3.10116 Value

**Fig 7.**
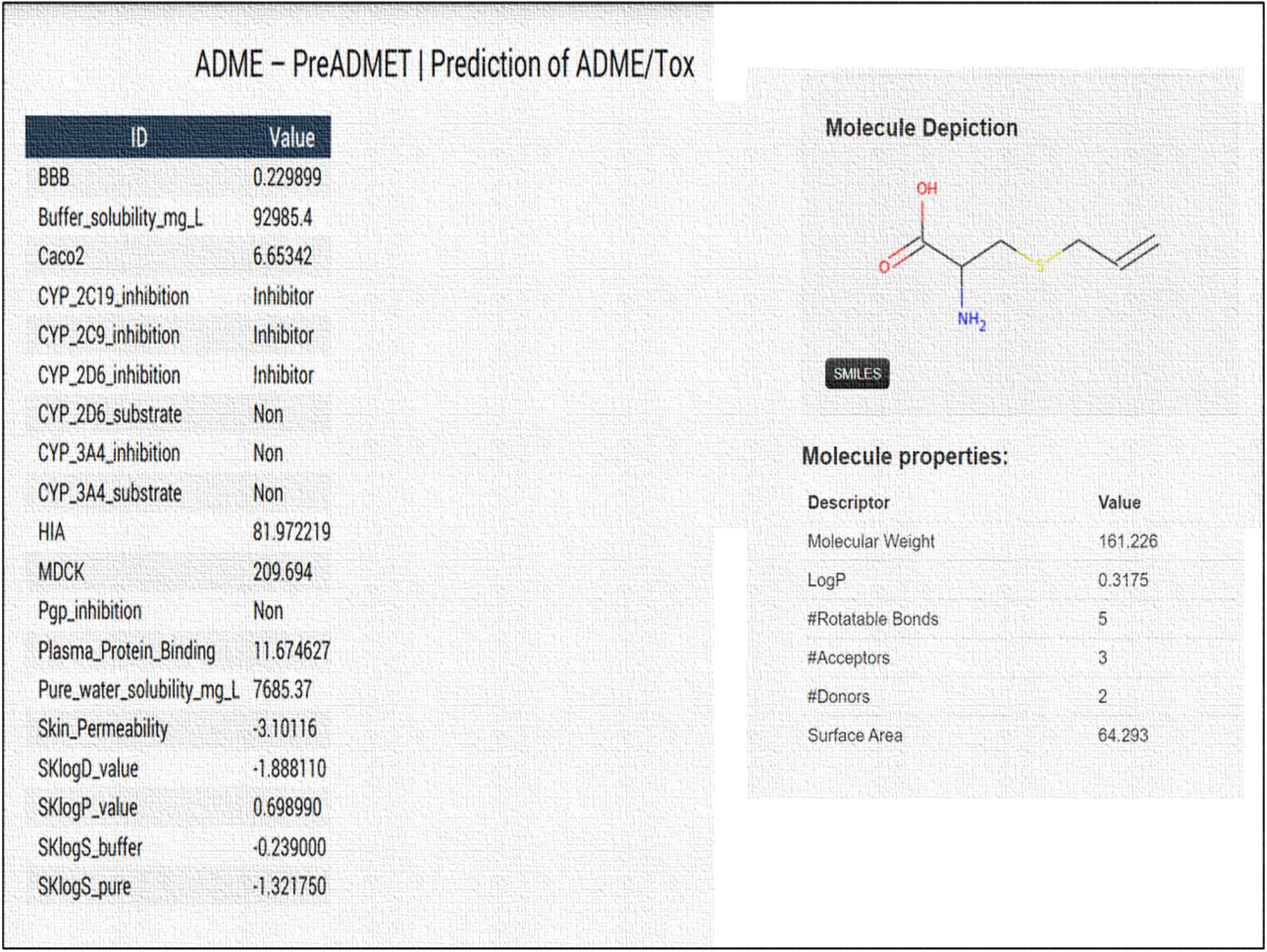
ADMET (metabolism, distribution, excretion, absorption, and toxicity) attributes of S-allylcysteine ( SAC ) predicted by PreADMET (https://preadmet.bmdrc.kr/)

In figure 8 we report ADMET (metabolism, distribution, excretion, absorption, and toxicity) attributes of S-allylmercaptocysteine (SAMC) predicted by PreADMET (https://preadmet.bmdrc.kr/ ):

- BBB (in vivo blood-brain barrier penetration (C.brain/C.blood) ≫ 0.190643 Value
- Calculated water solubility value in buffer system by SK atomic types (mg/L) ≫ 2537.48 Value
- in vitro Caco2 cell permeability (Human colorectal carcinoma; nm/sec) ≫ 1.60099 Value
- in vitro Cytochrome P450 2C19 inhibition ≫ Inhibitor
- in vitro Cytochrome P450 2C9 inhibition ≫ Inhibitor
- in vitro Cytochrome P450 2D6 inhibition ≫ Weakly
- in vitro Cytochrome P450 2D6 substrate ≫ No value
- in vitro Cytochrome P450 3A4 inhibition ≫ No value
- Human intestinal absorption (HIA, %) ≫ 82.576618 Value
- in vitro MDCK cell permeability (Mandin Darby Canine Kidney) ≫ 201.239
- in vitro P-glecoprotein inhibition≫ No value
- in vitro plasma protein binding (%) ≫ 0.000000
- in vitro skin permeability (transdermal delivery) ≫ −2.8397 Value

**Fig 8.**
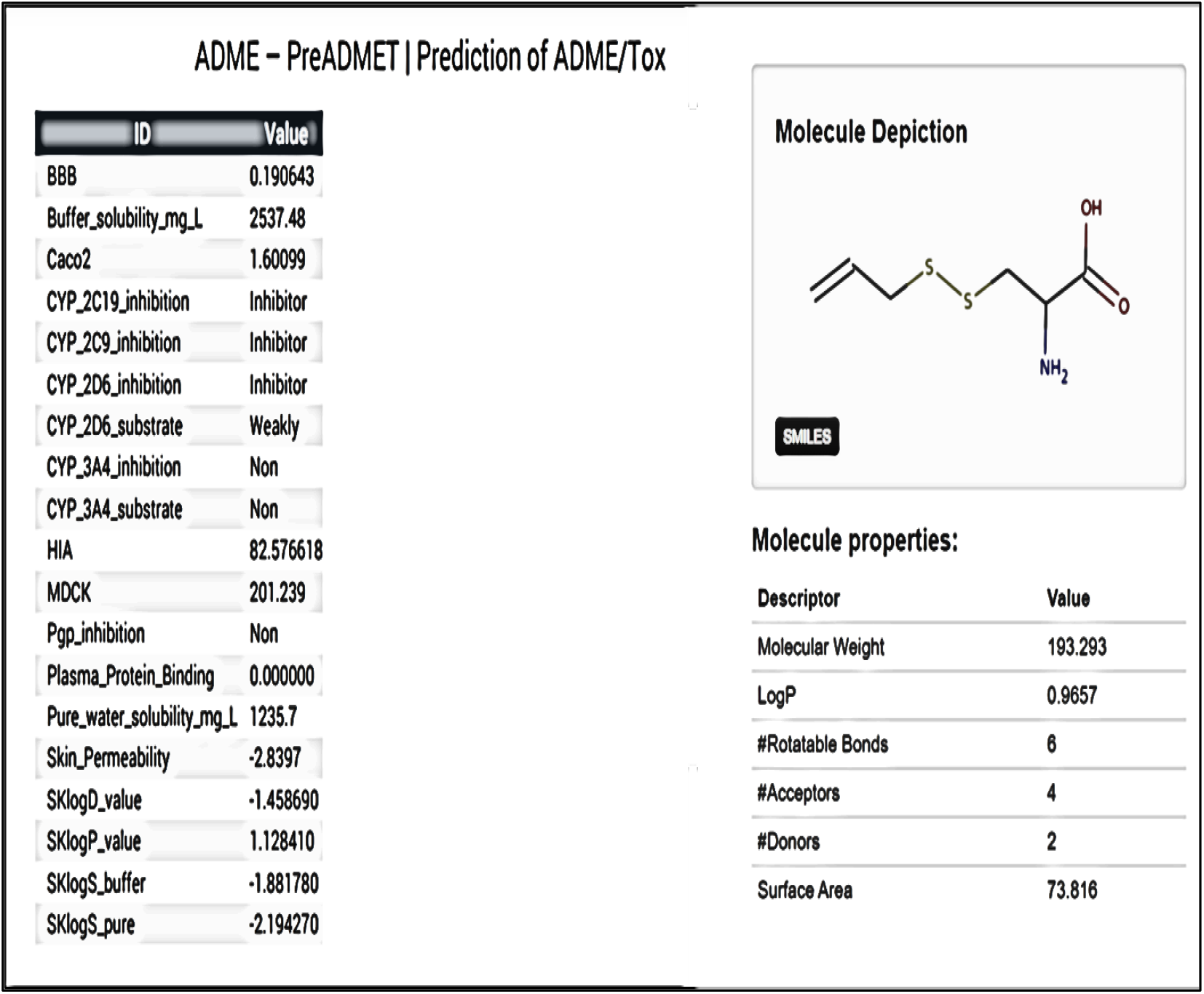
ADMET (metabolism, distribution, excretion, absorption, and toxicity) attributes of S-allylmercaptocysteine (SAMC) predicted by PreADMET (https://preadmet.bmdrc.kr/)[27]

#### 3.2.3 pkCSM (http://biosig.unimelb.edu.au/pkcsm/)

Drug development has a high attrition rate, with poor pharmacokinetic and safety properties a significant hurdle. Computational approaches may help minimize these risks It was seen that pkCSM performs as well or better across different pharmacokinetic properties than other freely available methods. This server is useful for Small-molecule pharmacokinetics prediction. In figure 9- 10 we report Prediction of pharmacokinetic properties: ADMET (metabolism, distribution, excretion, absorption, and toxicity) attributes of S-allylcysteine ( SAC ) and S-allylcysteine ( SAMC) predicted by pkCSM, respectively. (http://biosig.unimelb.edu.au/pkcsm/ [27]

**Fig 9.**
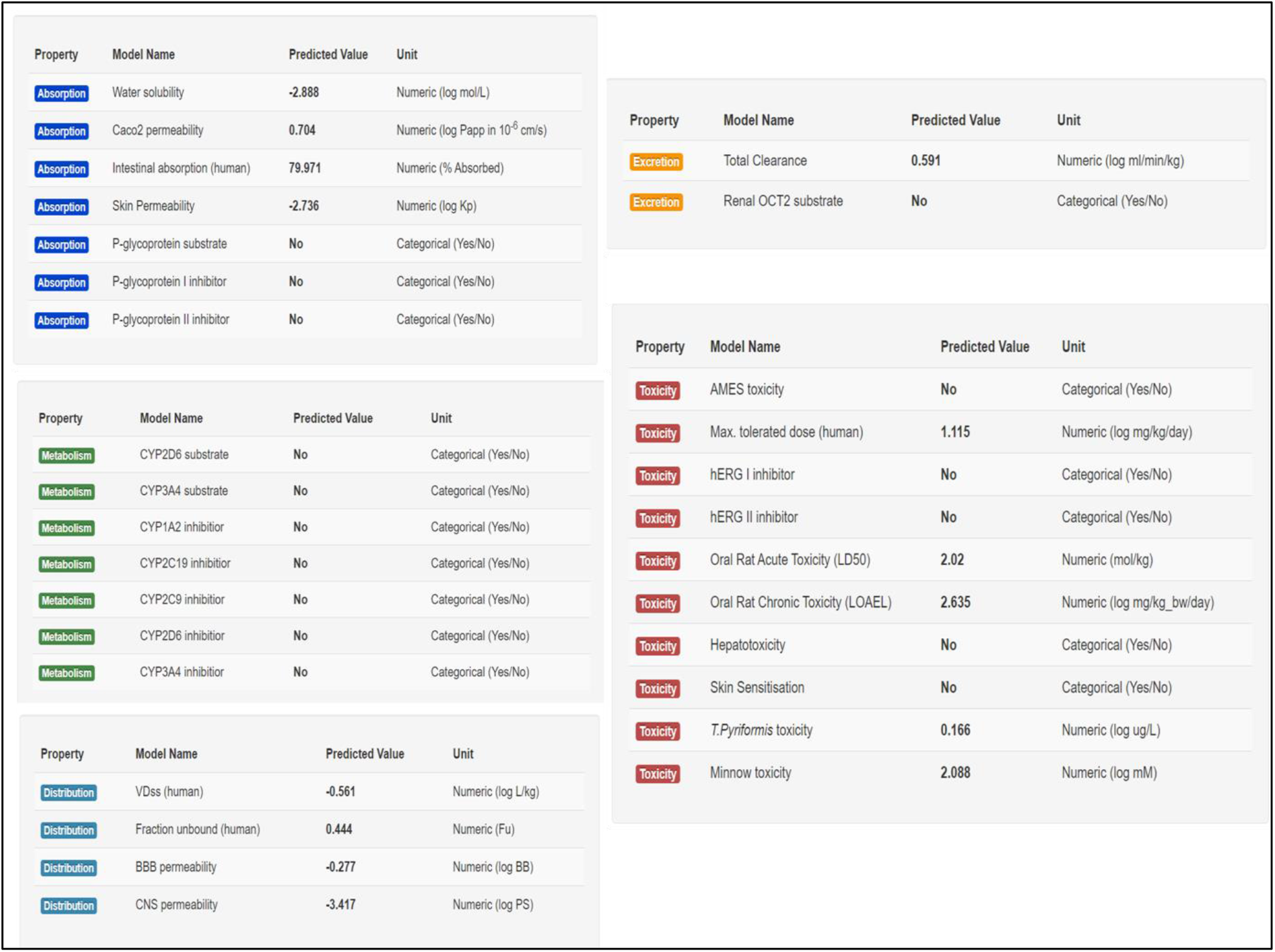
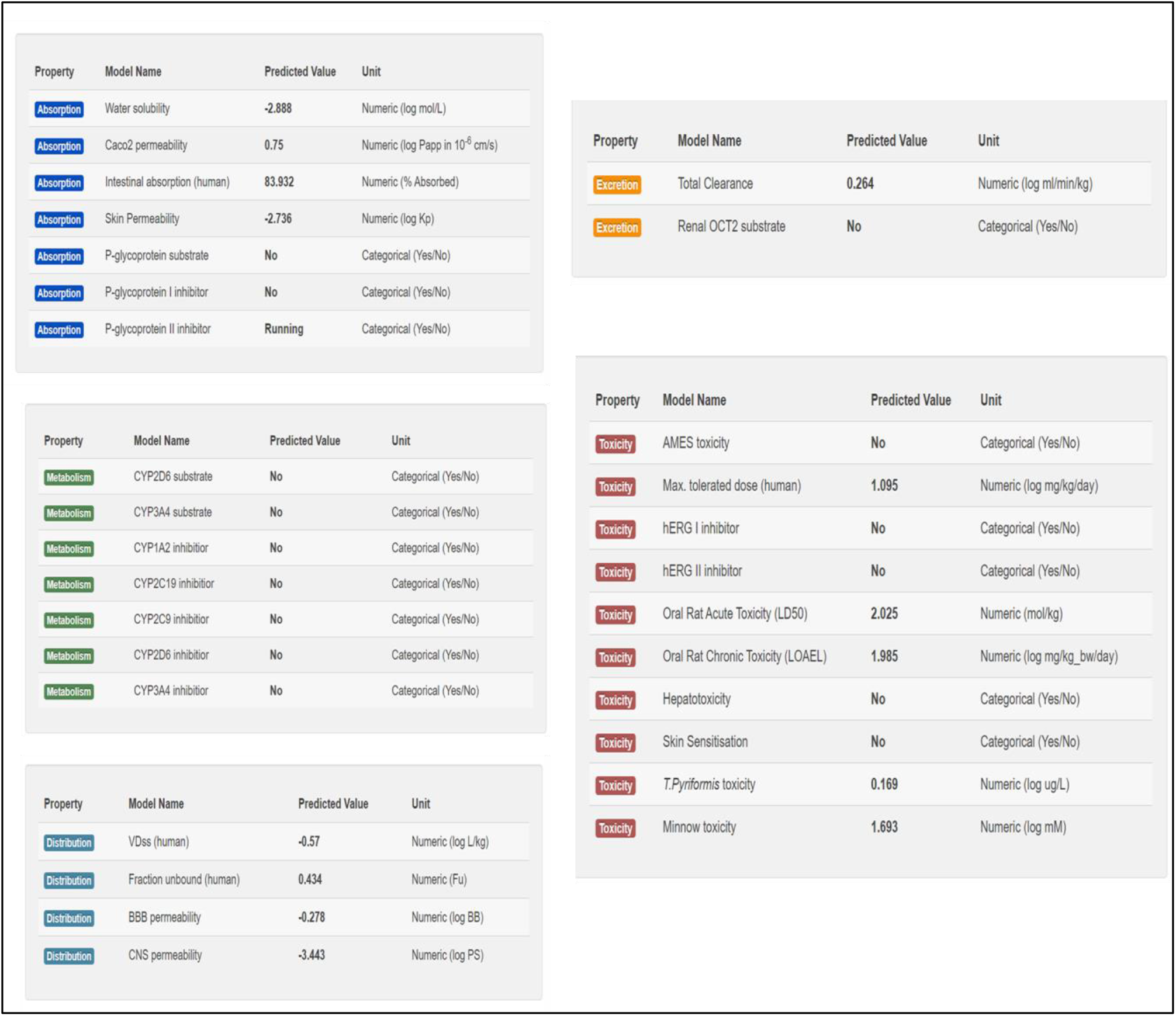
Prediction of pharmacokinetic properties: ADMET (metabolism, distribution, excretion, absorption, and toxicity) attributes of S-allylcysteine ( SAC ) SAMC) predicted by pkCSM (http://biosig.unimelb.edu.au/pkcsm/) [27]

## Conclusions

This paper encompasses the fundamental functions of open access in silico prediction tools, as PASS database (Prediction of Activity Spectra for Substances) that it estimates the probable biological activity profiles for compounds. This paper also aims to help support new researchers in the field of drug design and to investigate some of bioactive compounds in garlic. Particular attention we investigated Pharmacokinetic properties by several server, of S-allylcysteine ( SAC) and S-allylmercaptocysteine (SAMC) for their as anti-inflammatory, antibacterial, and antiviral, antioxidant, cardiovascular protective and anticancer property. Screening through each of pharmacokinetic criteria resulted in identification of Garlic compounds that adhere to all the ADMET properties. It was established an open-access database (PASS database, available bioinformatics tool SwissADME, PreADMET pkCSM database) servers were employed to determine the ADMET (metabolism, distribution, excretion, absorption, and toxicity) attributes of garlic molecules and to enable identification of promising molecules that follow ADMET properties. Further investigations will be conducted in Vitro and In vito by the SAMC, using Layered double hydroxides (LDH) ,which are one type of layered materials and are also known as anionic clays, are promising layered materials due to some of their interesting properties, such as to facile tunability of their composition, structure morphology and biocompatibility vehicle, against several cancer cells.

## Author contributions

I.V.F. conceived, designed and wrote the paper and performed the calculations and analyzed the data.

## Supporting Information

**Tab 2.**
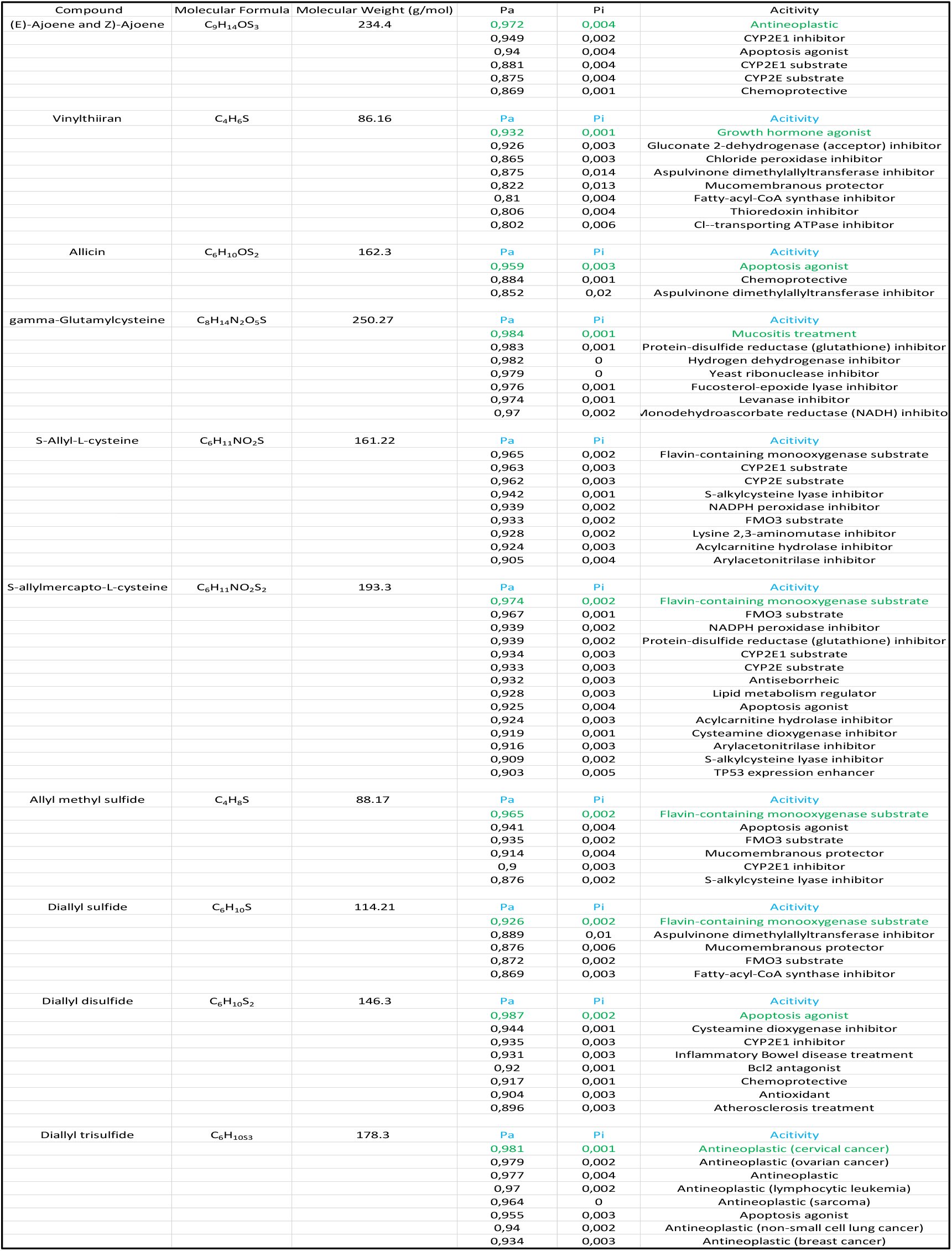
PASS Online Server (*Prediction of Activity Spectra for Substances) of Principal Organosulfur Compounds in Garlic . The output file represents a list of activities with two probabilities Pa (probability to be active) and Pi (probability to be inactive). Pa (probability “to be active“) estimates the chance that the studied compound is belonging to the sub-class of active compounds (resembles the structures of molecules, which are the most typical in a sub-set of “actives” in PASS training set). Pi (probability “to be inactive“) estimates the chance that the studied compound is belonging to the sub-class of inactive compounds (resembles the structures of molecules, which are the most typical in a sub-set of “inactives” in PASS training set). [8]*

**Tab 3.**
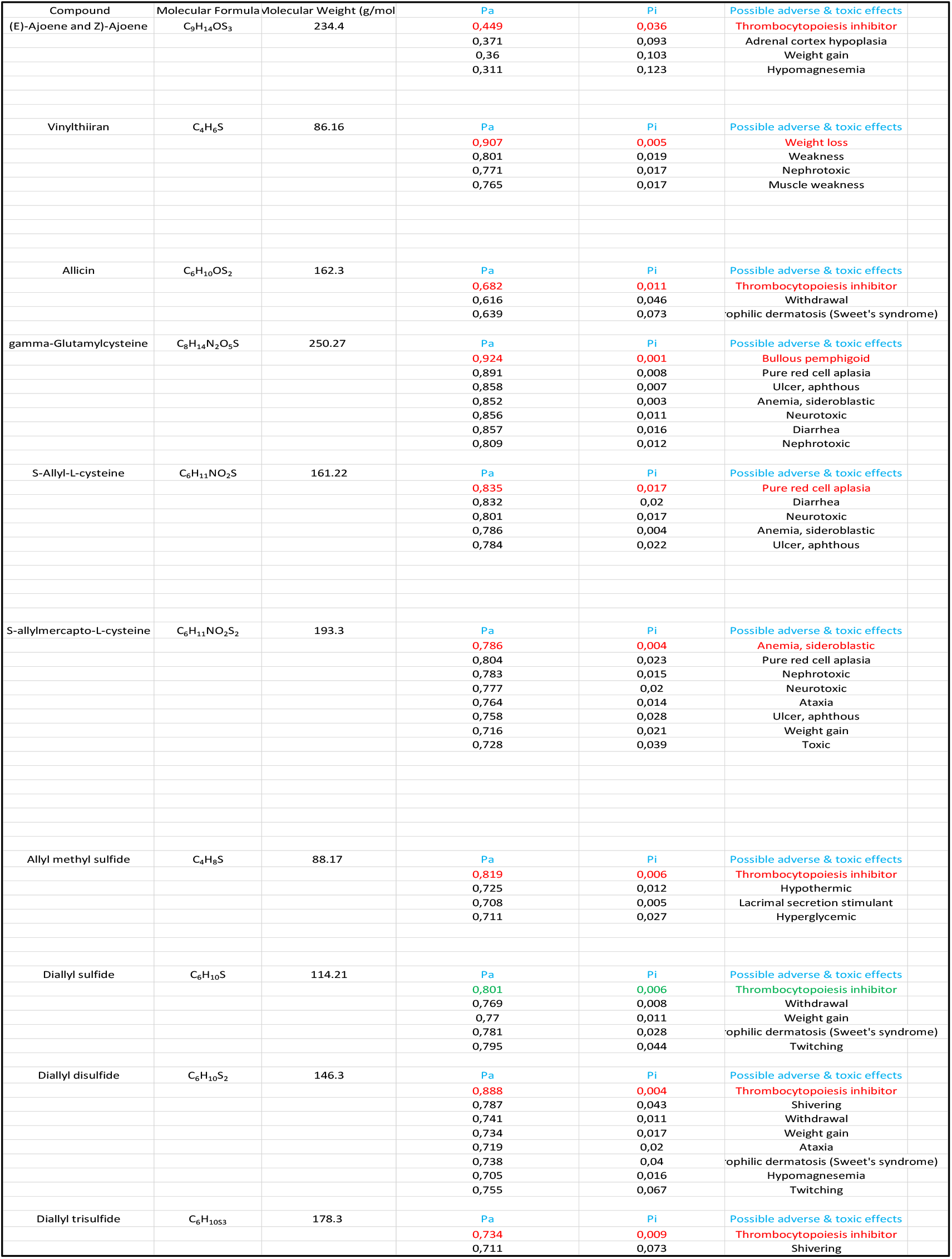
PASS Online Server (*Prediction of Activity Spectra for Substances) calculated possible adverse and toxic effects of Principal Organosulfur Compounds in Garlic [8]*

